# Characterization of Beta Tubulin Isotypes During Foam Cell Formation

**DOI:** 10.1101/141457

**Authors:** A. Torres, V. Contreras-Shannon

## Abstract

Foam cells contribute to the development of a cardiovascular condition called atherosclerosis. They arise when monocytes become engorged and lipid-laden after exposure to native low-density lipoproteins (Falk, 2006). It is assumed that the cytoskeleton is responsible for the morphological changes observed during foam cell formation. Beta tubulin and alpha tubulin are proteins that dimerize and polymerize to form microtubules, which are an important component of the cytoskeleton (Joshi, 1998). Little is known regarding the changes in cytoskeletal composition, particularly that of beta tubulins, throughout foam cell induction. The purpose of this study was to elucidate the expression patterns of beta tubulin isotypes 1–4 in human THP-1 monocytes throughout foam cell formation and to determine what relationship exists between beta tubulin expression and foam cell lipid aggregation. Levels of beta tubulin 1–4 were measured by western blot and immunofluorescence throughout the stages of foam cell differentiation, and beta tubulin isotypes were manipulated by siRNA to determine the effects of diminished beta tubulin expression on foam cell formation. Regardless of isotype, beta tubulin was always present in the highest amounts in monocytes. Levels of beta tubulin-1 and -4 were significantly decreased in macrophage and foam cells relative to monocytes (p < 0.0079, p < 0.0208, respectively). Beta 3 levels also exhibited a decrease. Beta 2 levels remained low regardless of differentiation stage. The distribution of beta tubulin 1 was shown to be more spindle-like (stretching across cells), compared to beta tubulins 2, 3, and 4, which exhibited a more “clumped,” less interconnected arrangement. When expression of beta tubulin isotypes 1, 3, and 4 were reduced in monocytes, resulting foam cells appeared to have more lipid aggregates and were significantly larger (p<0.0001) when compared to the size of foam cells without siRNA following treatment with PMA + LDL. In conclusion, the distribution of beta tubulins 1, 3, and 4 changes throughout the stages of foam cell induction, and manipulation of beta tubulins altered foam cell formation. Unexpectedly, the silencing or decreasing of beta tubulin enhanced lipid aggregation. Information concerning how beta tubulin expression can effect foam cell formation may offer insight into how to reduce plaque formation in patients with atherosclerosis.

## Introduction

A substantial portion of the normal function and movement of eukaryotic cells is mediated by the cytoskeleton, which has three well-established components: microfilaments composed of actin, intermediate filaments composed of various cell type-specific proteins, and microtubules composed of alpha and beta tubulin (Moseley, 2013). Arguably the most dynamic component of the cytoskeleton is the microtubule, which allows for functional changes in cellular architecture through the assembly and dissasembly of tubulin polymers (Joshi, 1998). Dynamic instability has been accepted as an inherent property of microtubules. Periods of rapid polymerization, where dimers of alpha and beta tubulin associate to form tubes of polarized rod-like polymers, are followed by periods of rapid depolymerization (Sept, 2007). Depolymerization is then either negated by another period of growth, or allowed to continue in order to mediate functions such as cell locomotion, chromosome separation during mitosis, and intracellular transport of organelles (Cooper, 2000).

Modulation of cell phenotype is particularly important when macrophages, the central effector cells of innate immunity, are involved (McWhorter, Wang, Nguyen, Chung, & Liu, 2013). Degree of morphological elongation of an activated macrophage has been associated with the macrophage’s orientation as either a pro-inflammatory M1 macrophage, or an effector cell of the pro-healing M2 orientation (McWhorter et al., 2013). The professional phagocytes, inclusive of macrophages, are most functionally adept at phagocytosis; the immediacy of the relation between the cytoskeleton and uptake of foreign bodies can be seen in the well-documented actin-dependence of phagocytic vesicle formation (Aderem & Underhill, 1999).

Of significance to this research is the involvement of macrophages in the pathogenesis of an immunoinflammatory disease of the cardiovascular system called atherosclerosis (Falk, 2006). Phagocytosis of foreign bodies occurs under normal physiological conditions, and is initiated by recognition of infectious agents through a variety of mechanisms including 1) activation of complement receptors, 2) activation of mannose receptors, and 3) antibody-mediated activation of Fc receptors (Aderem & Underhill, 1999). Activation of these receptors stimulates a change in the macrophage’s cytoskeleton to allow internalization of the detected antigen (Aderem & Underhill, 1999). Atherosclerotic lesions are most likely to occur where the endothelial layer of a vessel wall is defective or leaky, such as in the case of mechanical stress due to hypertension (Falk, 2006). Compromised endothelium is penetrable to circulating lipoproteins, which can be retained in the sub-endothelial space and undergo oxidation. Oxidized low-density lipoproteins (LDL) are chemotaxic and proinflammatory; activated endothelial cells overlaying the matrix containing trapped oxidized LDL secrete chemokines that prompt directional migration of blood-borne monocytes (Moore & Tabas, 2011). After diapedesis occurs, oxidized LDL is recognized by macrophage scavenger receptors including CD36 and Platelet-Activating Factor Receptor, which stimulates differentiation into the M2 phenotype (Rios et al., 2013). Atherogenic lipoproteins are then phagocytosed by activated macrophages causing the formation of foam cells, which are macrophages with high levels of intracellular lipid accumulation (Falk, 2006). Atherosclerotic plaques form over time with accumulation of foam cells in the subendothelial space of arteries (Falk, 2006). Intimal plaque can encroach on the vessel’s luminar space causing reduced blood flow, and in advanced cases, coronary artery thrombosis following plaque rupture (Falk, 2006).

Atherosclerosis is the most frequent underlying cause of coronary artery diseases, making it imperative to investigate how the formation of foam cells can be regulated. The current goals of atherosclerosis treatment include lowering the risk of clots forming, reducing lifestyle risk factors in order to delay or inhibit buildup of plaque, and preventing atherosclerotic-related diseases including coronary heart disease, carotid artery disease, and chronic kidney disease (“How Is Atherosclerosis Treated? - NHLBI, NIH,” n.d.). When dietary and lifestyle changes are not enough to control atherosclerotic progression, statins are often prescribed as a lipid-lowering therapy in order to reduce the risk of cardiovascular events (Cannon et al., 2004). In the event that modulation of beta tubulin levels has a significant effect on the formation of human foam cells, it is our hope that beta tubulin may be used as an additional disease target in the prevention or retardation of atherosclerosis progression. The modulation of microtubule dynamics by interactions with tubulin proteins have shown promise in the realm of anti-cancer therapy (Pasquier & Kavallaris, 2008). Through inhibition of the addition of tubulin molecules to microtubules, microtubule depolymerization can be induced (Hong, Chen, Donovan, Schneider, & Nisen, 1999). One such antimicrotubule drug, nocodazole, was shown in a previous study to inhibit the formation of foam cells in RAW 264.7 LDL-stimulated macrophages (Morishita et al., 2013). It was this finding that supported our hypothesis that there exists a relationship between beta tubulin expression and foam cell formation in a human-derived THP-1 monocyte cell line; such a relationship has not yet been elucidated.

In light of the cytoskeleton’s already-established role in phagocytosis and the maintenance of cellular architecture (Aderem & Underhill, 1999), the observation of morphological change during foam cell formation led to the following hypotheses: firstly, specific beta tubulin isotypes are involved in different stages of foam cell induction, and secondly, that disrupting specific beta tubulin isotypes in monocytes will alter LDL uptake and foam cell formation. This goal of this study was to determine the amount of each beta tubulin isotype 1–4 present in human monocytes at each stage in foam cell formation, and the effect beta tubulin isotype silencing had on monocyte differentiation into foam cells.

## Materials and Methods

### THP-1 Cell Line Maintenance, Culture, and Differentiation

THP-1 ATCC® TIB-202™cells were used in all experiments. For all plating and growth, RPMI 1640 medium with 4.5 g/L glucose, 10 mM HEPES, 1.5 g/L sodium bicarbonate, 1.0 mM sodium pyruvate, 10% fetal bovine serum, and 0.05 mM mercaptoethanol was used. For analyses of beta tubulin isotypes and their role in foam cell formation, typically 2 × 10^5^ cells were plated in multiwell dishes with or without coverslips. In the case of plasmid transfection, 1 x 10^5^ cells were plated in multiwell dishes. Cells were allowed to equilibrate in normal growth conditions described above for 24 hours prior to treatment or transfection. To induce macrophage and foam cell differentiation, THP-1 cells were treated with PMA (100 nM) or PMA (100 nM) + LDL (100 μg/mL), respectively, for 24 hours prior to analyses.

### Characterization of Beta Tubulin by Western Blot

To determine the amounts of each beta tubulin isotype (1–4) in monocytes (media-treated THP-1 cells), macrophages (PMA treated THP-1 cells), and foam cells (PMA + LDL treated THP-1 cells), we performed western blot. Cells were plated and differentiated according to the aforementioned specifications. For each of the four tubulins, cells in each treatment group (media only, PMA, and PMA +LDL) were prepared in triplicate. Lysates were prepared by adding 100 μL boiling 1X Laemmli sample buffer to wells after aspiration of media and washing of cells with PBS. Ten microliters of each lysate were loaded onto a 10% polyacrylamide gel. Each gel was run in duplicate. For immunodetection of β tubulins, blots were blocked with 5% milk with 0.1% Tween and then incubated in diluted primary mouse anti beta tubulin antibodies followed by secondary goat anti-mouse HRP-conjugated antibody (diluted 1:2500). Primary antibodies for beta tubulin −1, −3, and −4 were diluted 1:500, and antibody for beta tubulin-2 was diluted 1:250. (Beta-tubulin isotype-specific antibodies were a gift from Dr. Richard Ludueña.) Following immunodetection, blots were developed with chemiluminescent reagent and imaged using Carestream Image Station 4000MM Pro. After imaging, the same blots were washed thoroughly with PBS and reprobed with primary rabbit anti-GAPDH antibody (diluted 1:2500) and goat anti-rabbit HRP-conjugated secondary antibodies (1:2500); GAPDH was used as a loading control because the amount of protein loaded on the gel was not normalized beforehand. The ratio of the chemiluminescence of the tubulin bands to GAPDH bands was used to determine the relative levels of the beta-tubulin isotypes; Image J was used to quantify chemilumunescence.

### Elucidation of Isotype Distribution by Immunofluorescence

To determine isotype distribution in cells, immunofluorescence was used. Cells were plated according to aforementioned conditions. For each of the four beta tubulins, coverslips of PMA and PMA +LDL treated-cells were prepared in duplicate. Notably, immunofluorescence of media-treated monocytes was not performed as these cells do not adhere to coverslips and could not withstand the frequent washings and immunodetection steps. After 24 hour incubation in respective treatments, cells were fixed in neutral buffered formalin for a period of 10 minutes, permeabilized with ice-cold PBS+ 0.05% Triton X-100 detergent for 5 seconds, then blocked in 10% Normal Goat Serum. Following blocking, cells were incubated in primary mouse-anti beta tubulin antibodies diluted in Normal Goat Serum (1:100 for beta tubulin-1, -2, and -3, and 1:50 for beta tubulin-4) for 1 hour. Alexa Fluor 647-conjugated goat anti-mouse antibody (diluted 1:200) was applied following incubation with primary antibody. Following immunodetection, cells were stained with a 100 ug/mL Nile Red Stock Solution made in acetone and diluted 1:100 in PBS for 5 minutes. Coverslips were placed on microscope slides using hard-mount media containing DAPI, and imaged using a FLUOVIEW FV10i confocal microscope. To find appropriate laser settings for an isotype, preliminary scans across duplicates of media, PMA, and PMA + LDL-treated cells were performed, and the settings at which the highest beta tubulin signal could be detected without saturation was used for final microscopy of that isotype. Laser settings remained consistent across images taken within the respective isotype; i.e., all quantifications for beta tubulin-1 probed cells, regardless of treatment group, were imaged with the same laser settings. Other isotypes were imaged in the same manner.

Laser settings were not kept consistent across isotypes, as cross-isotype quantitative comparison was not performed.

### Silencing of β-Tubulin Expression With siRNA

Transfection with 25 picomoles of siRNA targeting beta tubulins −1, −3, and −4 was used to examine the effect of decreasing beta tubulin expression on foam cell formation. Human beta tubulin siRNAs were obtained from Santa Cruz Biotechnology: β1 Tubulin siRNA (sc-105000), β3 Tubulin siRNA (sc-105009), and β2C (β4) Tubulin siRNA (sc-105007). For the silencing study, cells for each treatment group were plated in triplicate. Following plating, a 10uM suspension of each siRNA was prepared, and cells were transfected with 25 picomoles siRNA using Santa Cruz Biotechnology transfection reagents, according to manufacturer’s instructions. Twenty-four hours post-transfection, induction of monocyte, macrophage, and foam cell formation was performed by addition of aforementioned treatments. Lysates were then prepared as previously described and blots were run in duplicate.

### Visualization of Lipids with Oil Red O

To confirm foam cell formation, cells were stained with Oil Red O (ORO). The ORO staining procedure began with fixation in neutral buffered formalin followed by staining with a solution of ORO (stock solution diluted 6:4 in nanopure water). Cells were imaged with a brightfield microscope.

### Overexpression of β Tubulin Expression with cDNA Plasmids

To examine the effect of increasing beta tubulin expression on foam cell formation, cells were transfected with plasmids encoding full length beta tubulin isotypes. Monocytes were plated as previously described, and then transfected with 500 ng of either beta tubulin isotype −1, −3, or −4 TrueClone full-length cDNA clones (OriGene) diluted in Opti-MEM (1x) Reduced Serum Medium. Lipofectamine® LTX with Plus™ Reagent (ThermoFisher) was added at the same time as the plasmids following the manufacturer’s procedure. After 24 hours, cells were treated with PMA and LDL as with previous experiments, and then subjected to Oil Red O and Western Blot procedures 24 hours later.

### Data Analysis and Statistics

All data is reported as mean +/-SEM. GraphPad prism was used to perform statistical analysis, specifically ANOVA and post-hoc tests (Bonferroni). ImageJ was used to analyze western blot images, micrographs for fluorescence measurement, and bright field microscopy for intracellular area measurements. For western blot, GAPDH and beta tubulin band intensities were measured and the resulting ratio was used to determine relative expression of beta tubulin. For immunofluorescence levels, corrected total cell fluorescence (CTCF) per unit area of a cell was calculated by multiplying the area of the selected cell by the mean fluorescence of the background readings for that microscopy picture, and subtracting that from the total fluorescence of the cellular beta tubulin contents. The CTCF value was then divided by the unit area to arrive at the final data value. Intracellular area of foam cells in micrographs was obtained by drawing freeform regions around foam cells.

## Results

### Characterization of Beta Tubulin by Western Blot

To determine the relative amounts of beta tubulin isotypes 1–4 throughout the different stages of foam cell formation, western blot was performed. Regardless of isotype, beta tubulin was always present in the highest amounts in monocytes (media-only treated cells) (Figure 1). Levels of beta tubulins-1 and −4 were significantly decreased in macrophage and foam cells relative to monocytes (p < 0.0079, p < 0.0208, respectively). Beta tubulin-3 levels also appeared decreased in macrophage and foam cells. Beta tubulin-2 levels remained about the same regardless of cell type. Given these findings, only beta tubulins −1, −3, and −4 were selected for silencing. *Elucidation of Isotype Distribution by Immunofluorescence*. To determine the spatial arrangement of each beta tubulin isotype *in vivo*, treated cells were probed with immunofluorescent antibodies for their respective tubulins. The distribution of beta tubulin-1 was shown to be more spindle-like (stretching across cells), compared to beta tubulins −2, −3, and −4, which exhibited a more “clumped,” less interconnected arrangement (Figure 2). Furthermore, quantification of beta-tubulin immunofluorescence confirmed that beta tubulin-1 and −3 immunofluorescence is slightly greater in foam cells compared to macrophage (Figure 3). Beta tubulin-2 immunofluorescence was slightly lower in foam cells when compared to macrophages.

**Figure 1:**
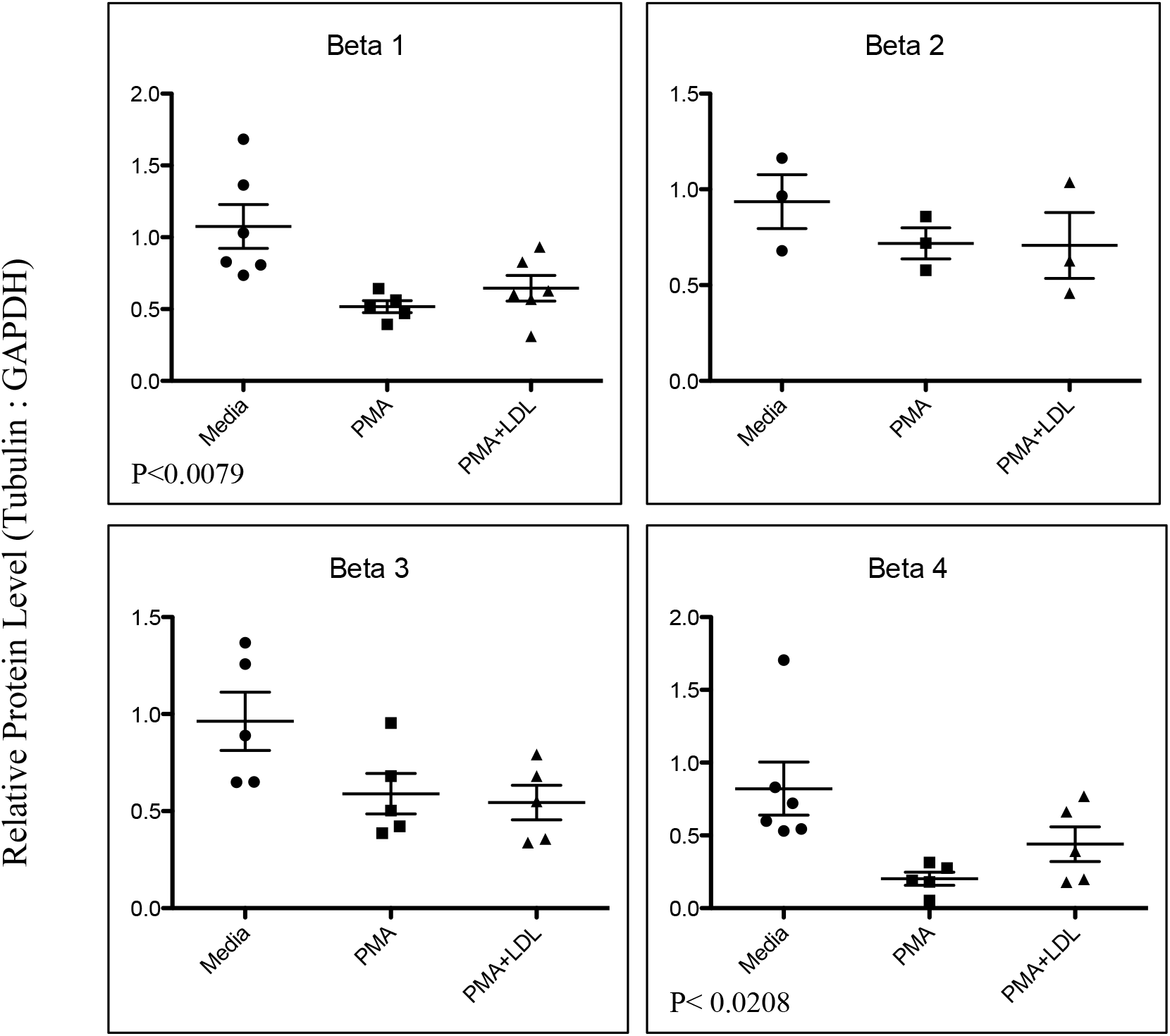
Relative Beta Tubulin Expression During Foam Cell Formation. Relative levels of beta tubulin expression were determined by treating cells with PMA (100 nM) and/or LDL (100 ug/ml) for 24 hours, preparing lysates, and performing western blot analysis for each isotype. For each graph, data points resulted from the blotting of 3 different lysates, with each lysate run in duplicate. Bars represent mean +/-SEM. One-way analysis of variance, p< 0.0079 (Beta 1) and p< 0.0208 (Beta 4).

**Figure 2:**
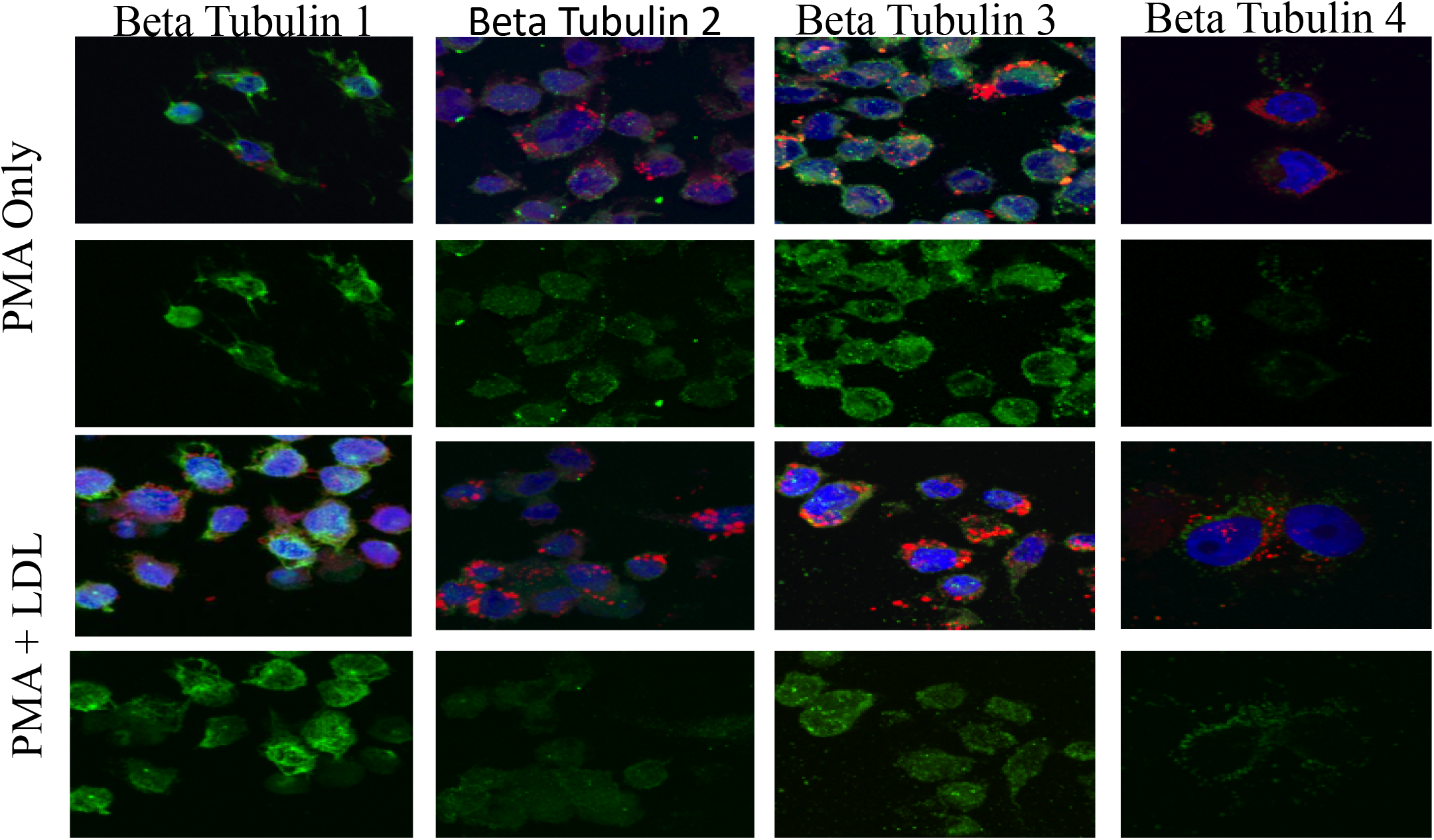
Immunofluorescence of Beta Tubulin. THP-1 cells treated with PMA (100nM), representing macrophages, or PMA (100 nM) and LDL (100 ug/ml), representing foam cells, were fixed, permeabilized, incubated in primary mouse anti-beta tubulin antibodies, then Alexa Fluor 647-conjugated goat anti-mouse antibodies, causing the respective beta tubulin isotype to fluoresce. In rows 1 and 3, tubulin appears green, the DAPI-stained nucleus appears blue, and Nile Red-stained lipids appear red. Rows 1 and 3 show all signals merged, while rows 2 and 4 exclusively feature the spatial arrangement of respective beta tubulins.

**Figure 3:**
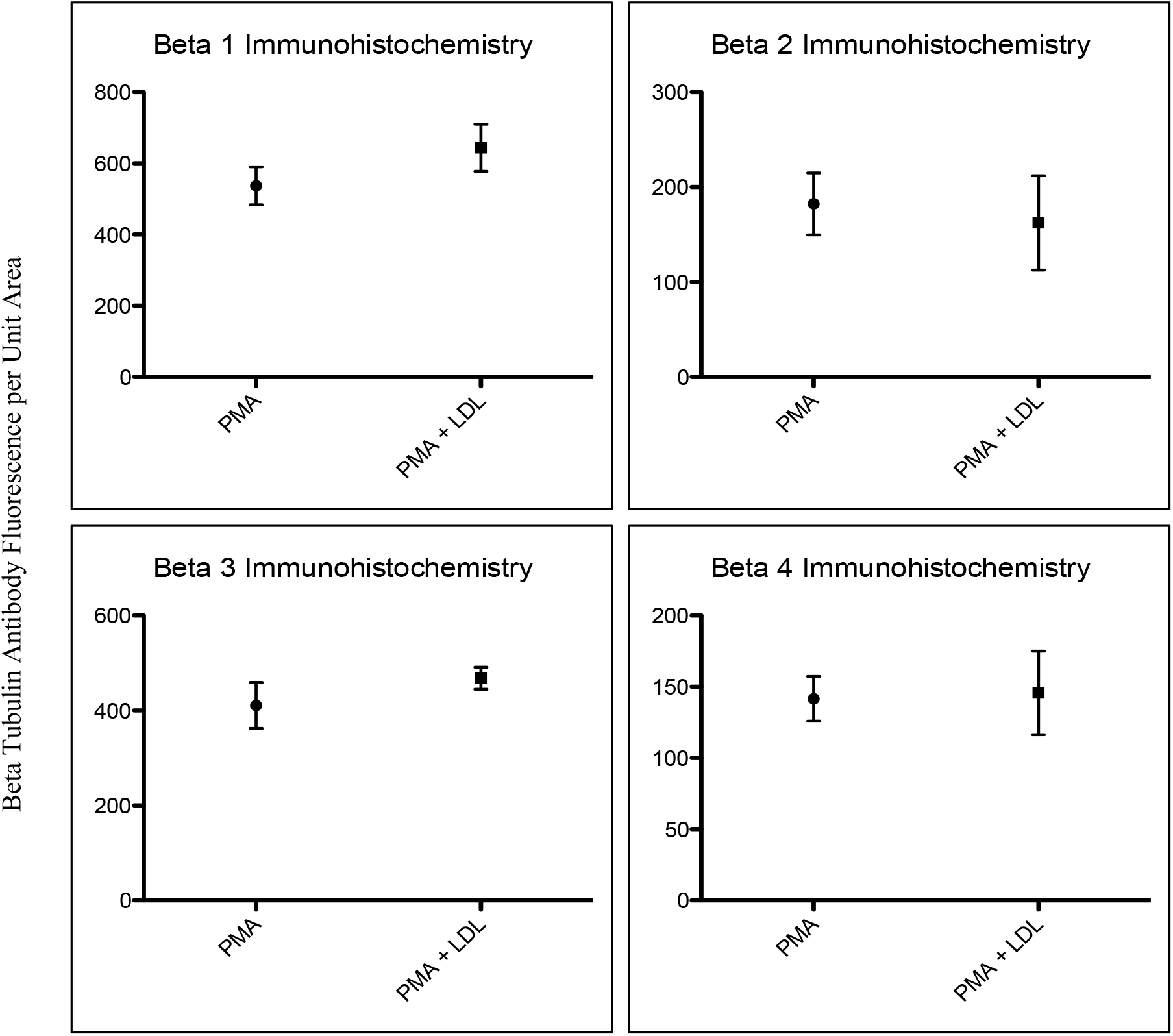
Relative Quantification of Beta Tubulin Immunofluorescence. THP-1 cells treated with PMA (100nM), representing macrophages, or PMA (100 nM) and LDL (100 ug/ml), representing foam cells, were fixed, permeabilized, incubated in primary mouse anti-beta tubulin antibodies, then Alexa Fluor 647-conjugated goat anti-mouse antibodies, causing the respective beta tubulin isotype to fluoresce. At least 9 cells per micrograph taken per group for each isotype were then analyzed by measuring the integrated density of the respective beta tubulin fluorescence signal, correcting for background signal, and dividing the corrected fluorescence by the unit area of the cell. On graphs, circles/squares represent mean, and bars represent +/-SEM.

### Silencing of Beta Tubulin Expression with siRNA

To determine the effects of manipulation of beta tubulin expression on a monocyte’s ability to undergo foam cell formation, we transfected THP-1 cells with siRNA to silence our beta tubulin isotypes of interest. First, we assessed the extent to which beta tubulins were silenced by siRNA. The beta-tubulin targeting siRNAs were successful in “knocking down” beta tubulin expression (row 1, Figure 4). All siRNA treated cells showed decreased levels of beta tubulin isotypes −1, −3, and −4 (p<0.0002, p<0.0001, p<0.0043 respectively). This was followed by evaluating foam cell formation after silencing beta tubulin expression. Silencing resulted in differences in lipid uptake. Cells with knocked-down beta tubulin-1, −3, and −4 showed a radical change in appearance (rows 2 and 3 of Figure 4). Cells appeared to have more lipid aggregates, and in general, exhibited a larger size when beta tubulin expression was less than normal. Cells with knocked down expression of beta tubulin −1, −3, and −4 were significantly larger (p <0.0001), when compared to the size of cells without siRNA following treatment with PMA and LDL (Figure 5). PMA-treated THP-1 cells with silencing of beta tubulin isotypes −1, −3, and −4 exhibited a 273.7%, 186.1%, and 281.4% increase in size, respectively, when compared to the control; when treated with PMA (100 nM) + LDL (100 ug/mL), silencing of beta tubulin isotypes −1, −3, and −4, resulted in 403.5%, 306.4%, and 389.3% increase in size, respectively, when compared to the control (Figure 5).

**Figure 4:**
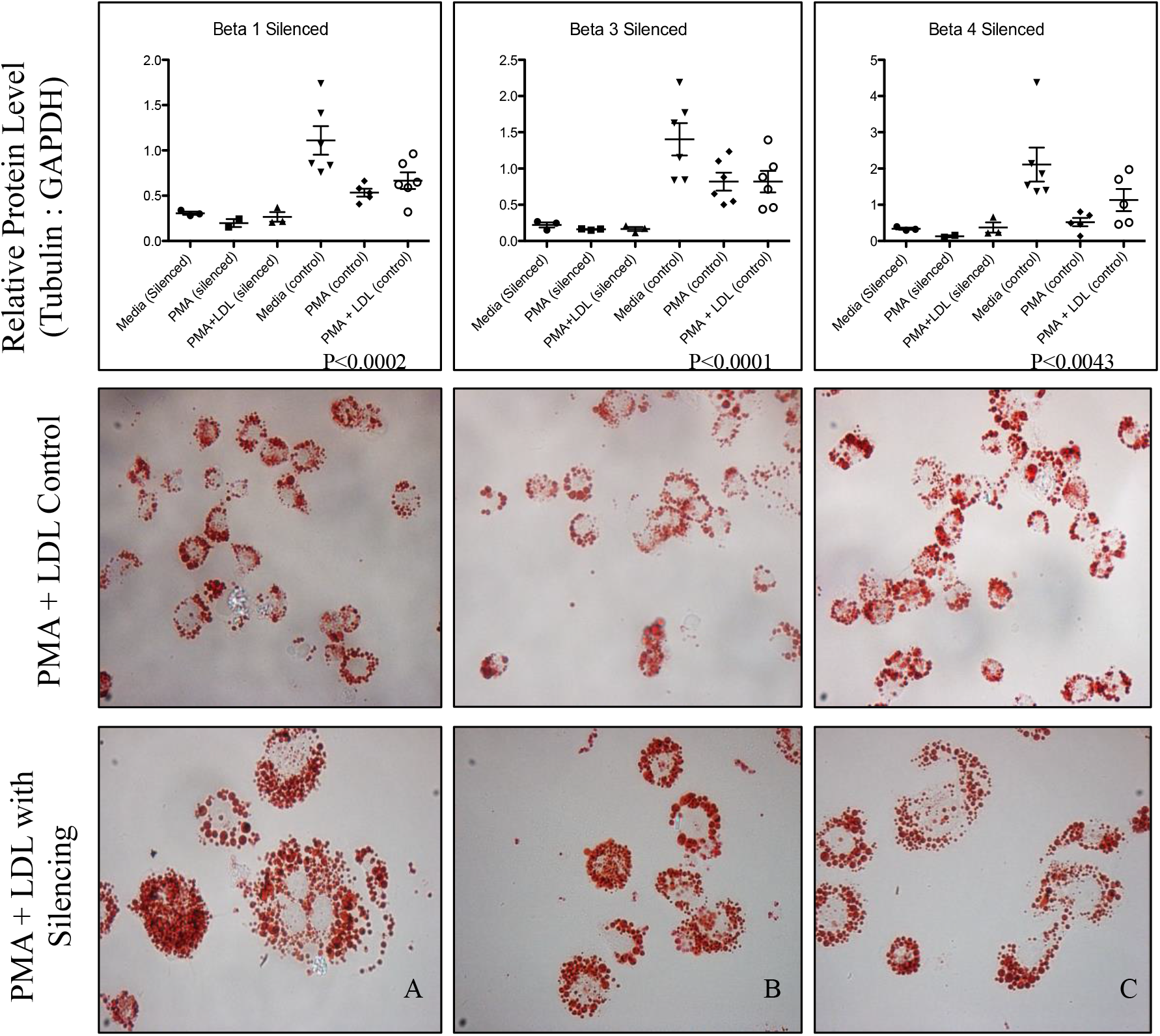
Beta Tubulin Protein Levels and Foam Cell Formation After Beta Tubulin Silencing. Graphs in top row show the beta tubulin:GAPDH ratio in siRNA-treated cells vs. normal cells. Micrographs in row two were cells treated with PMA and LDL but not siRNA. Micrographs in row three were cells transfected with 25 picomoles of siRNA using Santa Cruz Biotechnology transfection reagent for A) β1 tubulin, B) β3 tubulin, and C) β4 tubulin, left to incubate for 5 hours, then treated for 24 hours with PMA (100 nM) and LDL (100 ug/ml). Twenty-four hours after the addition of PMA and LDL, cells were stained with Oil Red O; red spots within cells are stained lipids. All cell microscopy was taken at a magnification of 400X.

**Figure 5:**
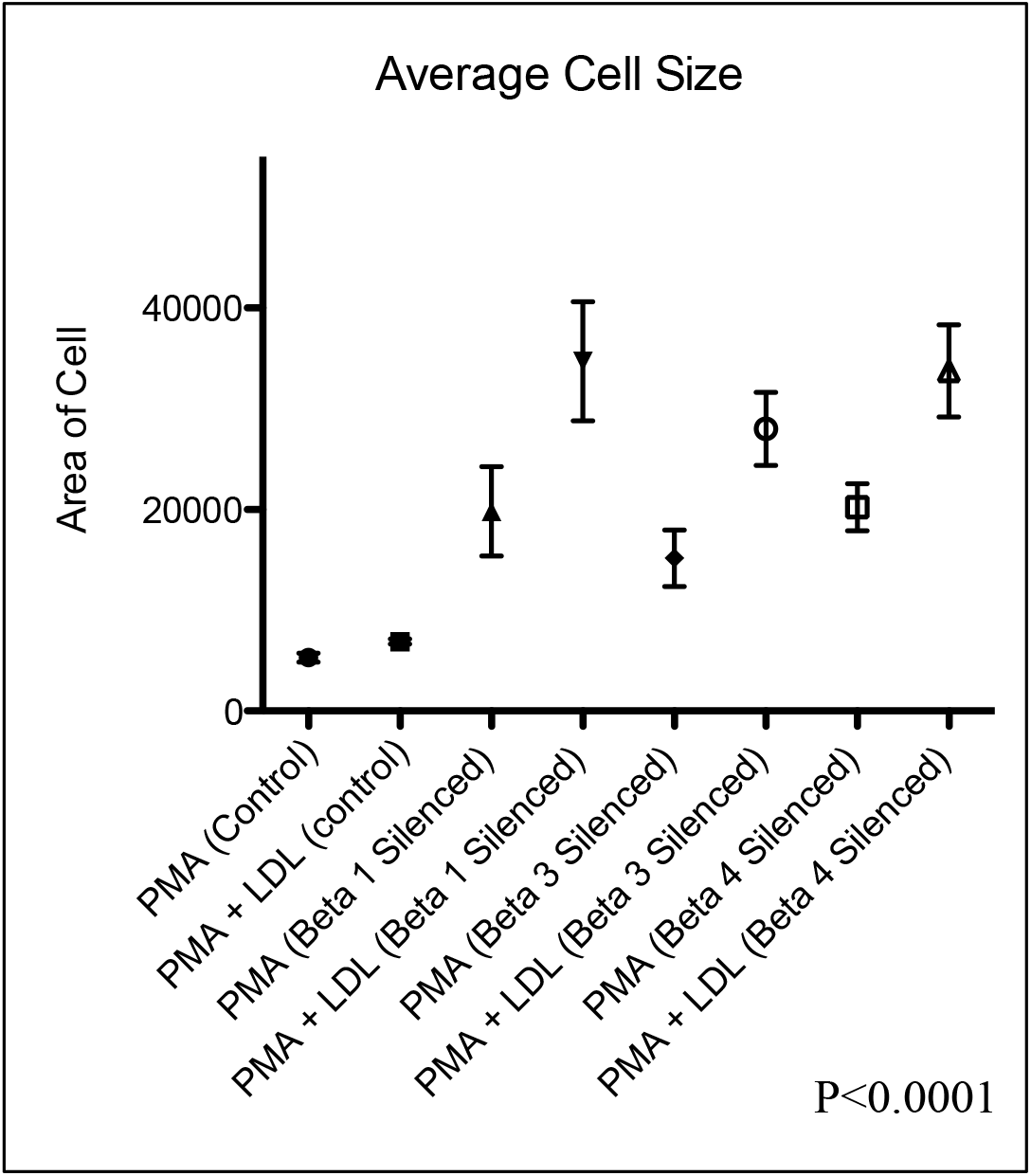
Average Size of PMA and PMA + LDL Treated THP-1 Cells After Silencing Beta Tubulins. THP-1 cells were transfected with 25 picomoles of siRNA using Santa Cruz Biotechnology transfection reagent and treated for 24 hours with either PMA (100 nM) or PMA and LDL (100 ug/ml). After 24 hours, cells were stained with Oil-Red O to facilitate microscopy. All microscopy pictures were taken at a magnification of 400X. At least 9 cells in each micrograph taken for each treatment group were then outlined in ImageJ, and evaluated for area. In graphs, shapes represent mean, and bars represent +/-SEM.

### Overexpression of Beta Tubulins with Full-Length cDNA Plasmids

To further elucidate the effects of manipulated beta tubulin expression on foam cell formation, we transfected cells with cDNA plasmids of the beta tubulin isotypes in question. First, we assessed the extent of beta tubulin overexpression with an overexpression plasmid. We were unable to quantify relative GAPDH:beta tubulin levels by western blot. Following transfection with respective overexpression plasmids, a change in the appearance of foam cells was not apparent, nor was there noticeable changes in lipid uptake between the control groups and groups transfected with beta tubulin plasmids (Figure 6). There was no significant size difference between the groups (Figure 7).

**Figure 6:**
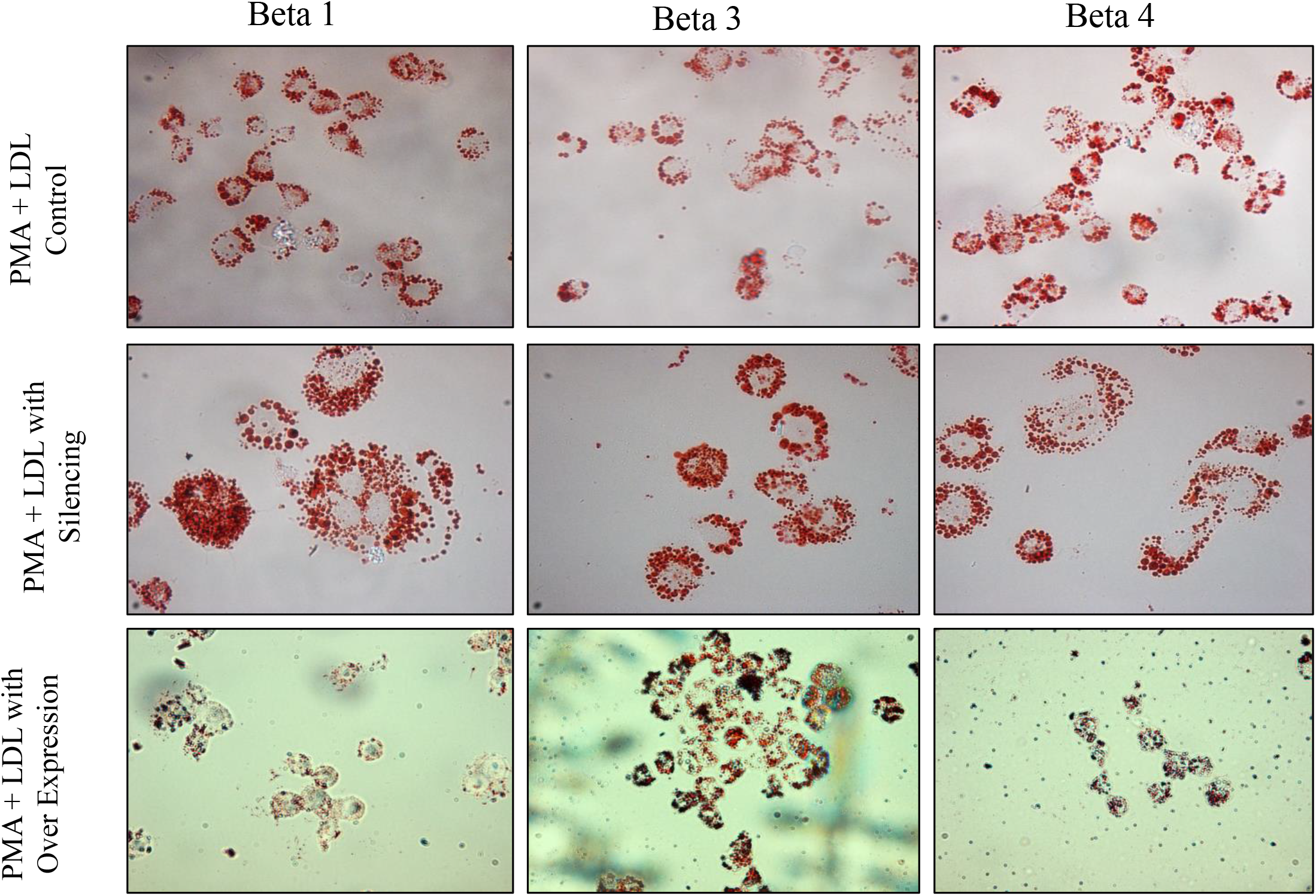
Comparison of Foam Cell Formation After Beta Tubulin Silencing and Suspected Beta Tubulin Overexpression. THP-1 Cells were treated with 24 hours with PMA (100 nM) and LDL (100 g/ml) following their respective transfections. The cells in row 2 were transfected with 25 picomoles of SiRNA using Santa Cruz Biotechnology transfection reagent for A) β1 Tubulin B) β3 Tubulin, and C) β4 Tubulin for five hours prior to incubation in PMA and LDL. Cells in row 3 were transfected with 500 ng of overexpression plasmid containing the full-length cDNA for A) β1 Tubulin B) β3 Tubulin, and C) β4 Tubulin for five hours before treatment with PMA and LDL. 24 hours after the addition of treatment, cells were stained with Oil Red O; red spots within cells are stained lipids. All cell microscopy was taken at a magnification of 400x.

**Figure 7:**
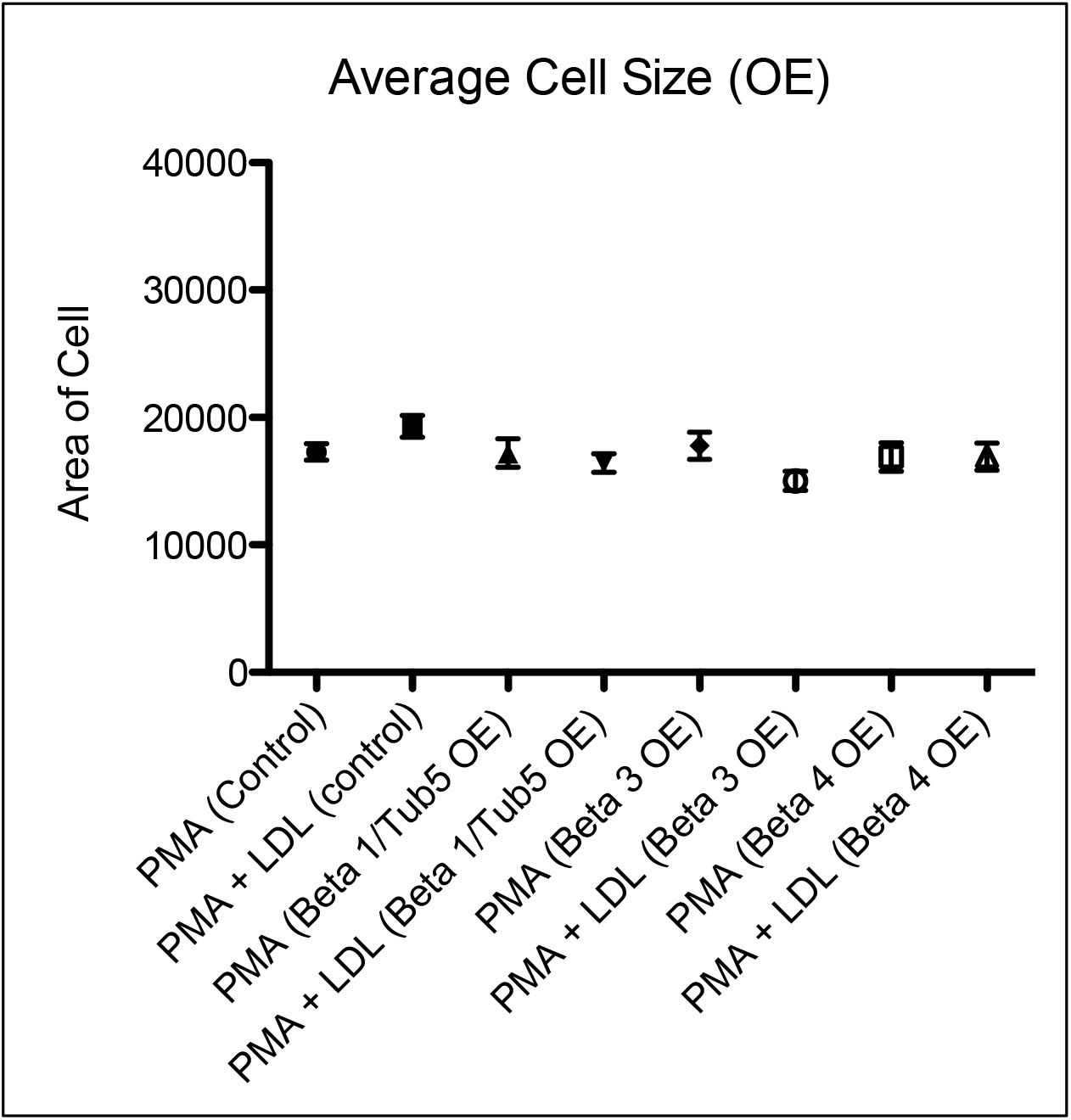
Average Size of PMA and PMA+LDL Treated THP-1 Cells With Suspected Overexpression (OE) of Beta Tubulin. THP-1 cells were transfected with 500 ng of overexpression plasmid containing the full-length cDNA for A) β1 Tubulin B) β3 Tubulin, and C) β4 Tubulin for before treatment with either PMA (100 nM) or PMA and LDL (100 ug/mL) for 24 hours. After 24 hours, cells were stained with Oil Red O to facilitate microscopy. All microscopy pictures were taken at a magnification of 400X. At least 9 cells in each micrograph per treatment group were then outlined in ImageJ, and evaluated for surface Area. In graphs, shapes represent mean, and bars represent +/-SEM.

## Discussion

The purpose of this study was to determine what relationship exists between four isotypes of beta tubulin found in humans and a human monocyte’s ability to undergo foam cell formation. These relationships were examined by 1) elucidating the amount of each beta tubulin isotype present in human monocytes at each stage in foam cell formation, and 2) observing the effect manipulation of expression of each isotype of interest has on monocyte differentiation. These findings suggest that 1) the distribution of beta tubulin isotypes −1, −3, and −4 changes throughout the three stages of foam cell induction, and 2) manipulation of beta tubulin expression alters lipid aggregation in foam cells. In light of the initial hypotheses, variations in beta tubulin expression at different points during foam cell formation were expected. However, the apparent relationship between amount of beta tubulin expression and both foam cell size and amount of lipid aggregation differed from our initial predictions.

During initial assessments of variations in native beta tubulin expression during foam cell formation, a spindle-like distribution of isotypes −1, −3, and −4 was observed. Distribution patterns coupled with a distinctive high-low-high expression pattern during monocyte, macrophage, and foam cell stages, respectively, suggested that these isotypes of beta tubulin are incorporated into microtubules used to control cell structure. We presumed that dramatic morphological changes in cell architecture, such as those observed during foam cell formation, were not likely to be directly contingent on the expression of a protein with a stable and unremarkable expression pattern. The distribution of beta tubulin-2 exhibited a more globular distribution than the other isotypes, and did not exhibit a significant change in expression throughout foam cell formation; it was for these reasons that beta tubulin-2 was not a candidate for manipulation of expression levels. The expression patterns of beta tubulin observed throughout foam cell formation, suggest mechanical rigidity, as it relates to microtubule quantity, would be least advantageous to leukocytes during periods generally associated with diapedesis and active or sustained phagocytosis. Leukocyte function is dependent on their ability to undergo deformation (Pierres et al., n.d.). The flexural rigidity of microtubules is increased as their stability is increased, and the rate of microtubule turnover is essential to mediating drastic changes in cell shape (Mickey & Howard, 1995). In light of recent research, it has been suggested that microtubules can influence the cellular symmetry and degree of pseudopodia projection in THP-1 cells; with the addition of microtubule-inhibiting drugs, exaggerated pseudopodia and asymmetric cell shape were observed, while the addition of microtubule-stabilizing drugs caused the cells to maintain a circular profile with reduced pseudopodia protrusions (Rosania & Swanson, 1996). For this reason, we inferred that the decrease in tubulin expression observed in THP-1 macrophages was consistent with their pseudopodia-mediated ability to undergo diapedesis and phagocytosis.

In cells with decreased expression of beta tubulin −1, −3, and −4, our findings demonstrate that both lipid aggregation and cell size are significantly altered compared to normal foam cells. This is in stark contrast to the findings of Morishita et al., whose experiments with the administration of the antimicrotubule drug nocodazole inhibited foam cell formation in mouse monoctyes exposed to similar conditions (Morishita et al., 2013). It was initially suspected that decreasing beta tubulin expression would in turn limit polymerization of microtubules, which would limit the size of the foam cell in order to maintain structural integrity. One possible explanation for the observed phenomena is that microtubules, in proper quantity, actually limit expansion of the cytoskeleton in the case of foam cell formation, rather than limiting cell size when in too paltry a quantity (Marshall et al., 2012). This is consistent with our previous observation that periods of active morphological expansions, characteristic of a macrophage’s prolonged phagocytic activity, are marked by periods of low expression of beta tubulin. It is possible that without the physical limitation of a microtubule-rich cytoskeleton, lipid aggregation and size expansion of foam cells went unchecked. Endocytotic vesicle formation, as in the case of lipid aggregation, does not appear to be inhibited by assumed hindrance of microtubule formation, while multi-nucleation of foam cells forged in conditions of low tubulin expression affirm that cell division is. Regardless, these findings clearly illustrate the importance of beta tubulin isotypes in regulating foam cell formation.

To confirm the inverse relationship between tubulin expression and the size and degree of lipid aggregation of foam cells, the trend must hold true in conditions of hyper-expression. Attempts were made to over-express beta tubulin isotypes −1, −3, and −4, and confirm elevated expression in relation to a to a housekeeping gene, GAPDH, by performing western blots and analyzing resulting band intensities. Due to the viscosity of lysates made from cells transfected with over-expression plasmids, we were unable to quantify relative expression, and therefore, we are not able to confidently assert that we overexpressed the protein of interest. It is suspected that the excessive amounts of DNA in the lysates due to the addition of plasmids made running the western blot unsuccessful. Ongoing attempts are being made to circumvent this problem so that the aforementioned relationship can be better elucidated. We expect that in the definitive case of successful overexpression, foam cell size and lipid content of experimental foam cells will either closely mirror that of normal foam cells, or exhibit a smaller, less lipid-laden appearance.

Given these findings, that a there exists a relationship between beta tubulin expression and the size and degree of lipid aggregation within foam cells, we will identify a point of regulation in the pathogenesis of atherosclerosis. In the event that targeted treatment can prevent foam cells from forming, the progression of atheroscerlotic plaques may be delayed, and the risk of rupture and coronary thrombosis that account for the mortality and morbidity associated with atherosclerotic disease may be minimized (Falk, 2006).

